# Fission and PINK-1-mediated mitophagy are required for Insulin/IGF-1 signaling mutant reproductive longevity

**DOI:** 10.1101/2021.08.16.456566

**Authors:** Vanessa Cota, Coleen T. Murphy

## Abstract

Women’s reproductive cessation is the earliest sign of human aging and is caused by decreasing oocyte quality. Similarly, *C. elegans’* reproduction declines with age and is caused by oocyte quality decline. Aberrant mitochondrial dynamics are a hallmark of age-related dysfunction, but the role of mitochondrial morphology in reproductive aging is largely unknown. We examined the requirements for mitochondrial fusion and fission in oocytes of both wild-type worms and the long-lived, long-reproducing insulin-like receptor mutant *daf-2*. We find that normal reproduction requires both fusion and fission. By contrast, *daf-2* mutants require fission, but not fusion, for reproductive span extension. *daf-2* mutant oocytes’ mitochondria are punctate (fissioned) and may be primed for mitophagy, as loss of the mitophagy regulator PINK-1 shortens *daf-2’s* reproductive span. Our data suggest that *daf-2* maintain oocyte mitochondria quality with age via a shift toward punctate mitochondrial morphology and mitophagy to extend reproductive longevity.

## Introduction

Mitochondria are dynamic organelles that undergo fission and fusion throughout their lifetimes in response to energetic cues, damage, and needs of the cell. Fission, which is controlled by the conserved dynamin-related protein DRP-1, separates elongated mitochondria into smaller, punctate mitochondria (Labrousse et al., 1999; Smirnova et al., 2001). Fusion is the opposing process by which mitochondria form elongated, tubular structures, and is controlled by transmembrane GTPases Mfn1 and Mfn2 in mammals, and FZO-1 in yeast and *C. elegans* (Hales and Fuller, 1997; Santel and Fuller, 2001; van der Bliek et al., 2013). The balance of fission and fusion is necessary to maintain the health of the mitochondria population in the cell; healthy mitochondria are maintained by either compensating for damaged proteins (fusion), or by eliminating damaged proteins (fission and subsequent mitophagy). A disruption in dynamics is a hallmark of mitochondrial dysfunction, a common sign of aging and age-related disease (Srivastava, 2017).

Cessation of female reproduction is the first human age-related decline, and has become increasingly important as the average maternal age increases (Martin et al., 2013; Mills et al., 2011). The main contributor to reproductive decline is the age-associated decline of oocyte quality, rather than an overall loss of oocytes (te Velde, 2002). The mitochondrion is a prime candidate for influencing reproductive aging: mitochondria are the most abundant organelle in the mature oocyte (May-Panloup et al., 2007), and mitochondrial dysfunction has been associated with reproductive aging and human oocyte quality decline (Kasapoğlu and Seli, 2020; Zhang et al., 2017). Mitochondrial dynamics may also play a role in reproductive health. Mouse oocytes with reduced levels of Mitofusins Mfn1 and Mfn2 show signs of mitochondrial dysfunction, coupled with reproductive dysfunction (Liu et al., 2016; Zhang et al., 2019). Levels of Drp1 activity are reduced in aged oocytes, and Drp1 KO mice have reduced fertility due to fewer oocytes reaching maturity (Udagawa et al., 2014). These data support a role for fission and fusion in reproduction, but many unanswered questions remain; specifically, how fission and fusion factors affect reproductive decline with age, and whether specific mitochondrial dynamics promote reproductive longevity.

*C. elegans* are a useful model to study reproductive aging. Like humans, *C. elegans’* reproduction declines with age. At the end of reproduction, wild-type *C. elegans* germ cells (Garigan et al., 2002) and oocytes (Luo et al., 2010) deteriorate, fewer oocytes are fertilized, and fewer fertilized embryos hatch (Hughes et al., 2005; Luo et al., 2010). Therefore, like humans, it is the decline of *C. elegans* oocyte quality, not the number of oocytes, that determines reproductive span. Furthermore, mutants that extend reproductive span have been identified, including the insulin/IGF-1 signaling (IIS) receptor mutant *daf-2* (Huang et al., 2004). While longevity and reproductive span extension can be uncoupled, such as in TGF-beta mutants (Luo et al., 2009), insulin/IGF-1 receptor mutants (Kenyon et al., 1993) are long-lived and have a long reproductive span (Huang et al., 2004) due to the activation of the downstream FOXO transcription factor DAF-16 (Luo et al., 2009). *daf-2* mutants maintain germ cell integrity with age (Garigan et al., 2002), have fewer unfertilized oocytes, and produce more embryos reaching maturity - all markers of high-quality oocytes (Luo et al., 2010; Templeman et al., 2018).

Studies of mitochondria in mammalian and *C. elegans* somatic tissues have led to the prevailing model that the elongated morphology is associated with “youthfulness,” while punctate (“fragmented”) mitochondria are a sign of dysfunction, a hallmark of aging and age-related disease (Srivastava, 2017). Mitochondria in *C. elegans* muscle cells (Lamarche et al., 2018; Palikaras et al., 2015; Regmi et al., 2014; Wang et al., 2019; Weir et al., 2017) and neurons (Jiang et al., 2015; Morsci et al., 2016) are elongated in youth and fragment with age. *daf-2* and mutants of the IIS pathway maintain the elongated mitochondria structure in body wall muscle and neurons longer than does wild type with age (Jiang et al., 2015; Lamarche et al., 2018; Morsci et al., 2016; Regmi et al., 2014; Wang et al., 2019). These findings support a model in which mitochondrial elongation (fusion) is indicative of healthy, functional mitochondria in somatic tissue.

Here we investigated the relationship between mitochondrial morphology and reproductive aging. In contrast to earlier models of mitochondrial health, we find that the oocytes of the reproductive longevity mutant *daf-2* maintain punctate mitochondrial morphology throughout reproduction, while wild-type oocyte mitochondria maintain their elongated morphology with age, even while losing reproductive capacity. Therefore, elongated mitochondrial morphology is uncoupled from youthful function in both *daf-2* and wild-type oocytes. We also find that wild-type oocytes require both fission and fusion for reproductive health, but *daf-2* requires only fission (not fusion) as well as PINK-1-mediated mitophagy to maintain oocyte quality and reproductive longevity. Taken together, our data support an alternative strategy for oocyte quality maintenance that engages mitochondrial dynamics and elimination to promote reproductive longevity.

## Results

### *daf-2* oocytes display punctate mitochondria

The insulin-like receptor mutant *daf-2* has both an extended lifespan (Kenyon, et al. 1993) and an extended reproductive span (Figure 1A; Huang et al., 2004; Luo et al., 2009). *daf-2* maintains its extended reproductive span by maintaining oocyte quality with age (Luo et al., 2010; Templeman et al., 2018). Further, in an additional test of reproductive capability, aged hermaphrodites are mated with young males that produce wild-type sperm to determine the fraction of the population able to produce progeny with age. We mated aged (Day 7) wild-type and *daf-2* hermaphrodites and found that significantly more *daf-2(e1370)* were still able to produce progeny (Figure 1B). By Day 8 of adulthood, *daf-2(e1370)* mutants still maintain large, uniform, cuboidal oocytes (Luo et al., 2010), Figure 1C), while wild-type oocytes deteriorate (Figure 1C-D; (Luo et al., 2010)).

**Figure 1.**
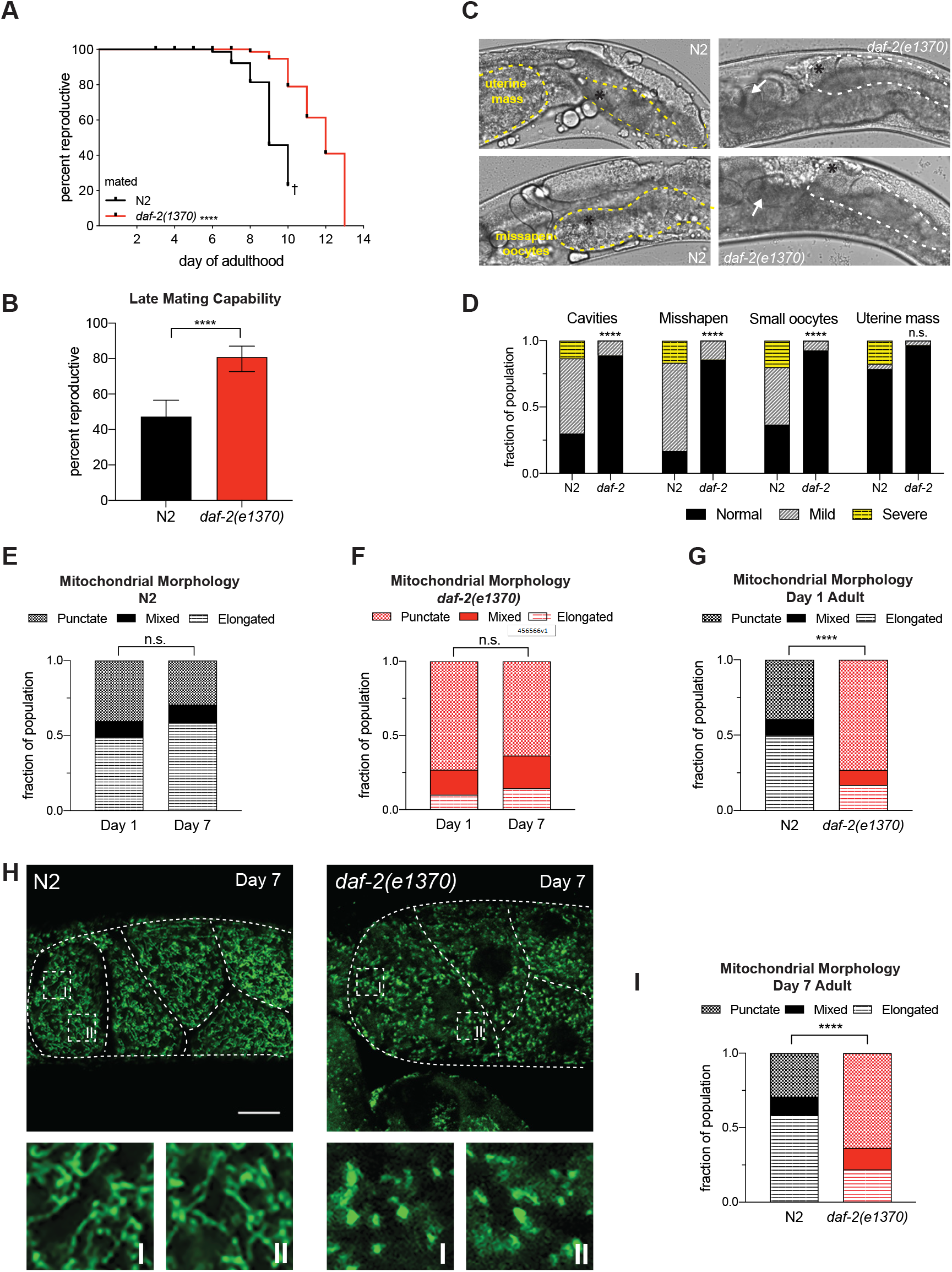
*daf-2* oocytes display punctate mitochondria *daf-2(e1370)* has an extended reproductive span (Huang, et al. 2004) in hermaphrodites (n=96-121) mated with males (Luo et al. 2010). † indicates loss of hermaphrodites due to high rates of matricide by bagging. (B) *daf-2(e1370)* late-mating capability is increased (n=115), as compared to N2 (n=110), when males were provided to aged Day 7 adult hermaphrodites for mating. Error bars represent 95% confidence interval. (C) On Day 8, N2 have deteriorated germlines (yellow dotted outline), while *daf-2(e1370)* retain cuboidal shaped oocytes (white dotted outline). Black asterisks mark the −1 oocyte, white arrows highlight developing embryos. Scale bars 20um. (D) *daf-2(e1370)* maintains oocyte quality (Luo et al. 2010) in mated Day 8 hermaphrodites (n=27-30). (E-H) Quantification of mitochondria morphology in the −1 oocyte. Germlines were dissected, fixed, and stained with anti-ATP5α to mark mitochondrial membranes. Each Z-stack was blindly-scored into 3 categories: punctate, mixed, or elongated. Mitochondria morphology does not change with age in N2 (E) and *daf-2(e1370)* (F) (in E and F, images were collected and blindly scored against the opposing genotype at each timepoint and the morphology ratios were compared against each other). (G) In Day 1 adults N2 (n=117) have majority elongated mitochondrial morphology while *daf-2(e1370)* have majority punctate morphology (n=107). (H) Images of mitochondrial morphology in Day 7 adults in N2 (left, elongated), and *daf-2(e1370)* (right, punctate). Scale bar 20um. White-dotted boxes are enlarged sections of the −1 oocyte (below). Images in H were processed using the same parameters across the entire image (i.e. adjusting for brightness and contrast). (I) Quantification of morphology in Day 7 adults. In Day 7 adults N2 −1 oocytes have majority elongated mitochondrial morphology (n=83). *daf-2(e1370)* oocytes have majority punctate morphology (n=118). Three independent replicates were imaged, scored, and pooled together for graphs E-I. ****p ≤ 0.0001.

To determine how mitochondria impact oocyte quality and reproductive longevity, we first examined mitochondrial morphology. In muscle cells, *daf-2* prevents mitochondrial fragmentation during aging (Chaudhari and Kipreos, 2017; Regmi et al., 2014; Wang et al., 2019) and neurons (Jiang et al., 2015; Morsci et al., 2016), therefore, we hypothesized that *daf-2* may also prevent oocyte mitochondrial fragmentation and maintain elongated mitochondria with age. To test this hypothesis, we examined the mitochondrial morphology of the most mature oocyte, the −1 oocyte, in Day 1 and Day 7 wild-type and *daf-2(e1370)* adults. Using a mitochondrial membrane marker (ATP5α), oocytes were imaged and blindly scored for three categories characterizing morphology. (Blinded, manual scoring was chosen instead of measuring mitochondria length or perimeter/area, strategies sometimes employed in flat muscle cells, because 1) the −1 oocyte is a large three-dimensional cell that does not organize its mitochondria on a single plane and 2) mitochondria are densely packed into oocytes, making it difficult to discern the beginning and end of each mitochondrion when elongated.) First, we found that wild-type oocyte mitochondria appear elongated and tubular in young Day 1 adults and that this morphology does not change with age (Day 7 of adulthood; Figure 1E and Supplemental Figure S1A). This result suggests that the elongated state of mitochondria in wild-type oocytes does not correlate with “health,” as this morphology persists even as the oocytes lose function.

Because high-quality mitochondria are often described as “elongated” while low-quality mitochondria are described as “fragmented,“ we were surprised to find that the mitochondria in *daf-2* −1 oocytes are not primarily elongated and tubular; instead, *daf-2* oocyte mitochondria appear punctate in both young (Day 1) and old (Day 7) animals (Figure 1F and Supplemental Figure S1B), suggesting that rather than fragmenting from an originally elongated state, *daf-2* oocyte mitochondria maintain a punctate state throughout their lives. Furthermore, at Day 1 (Figure 1G) and Day 7 (Figure 1H and 1I, Supplementary Videos 1-6) *daf-2* has significantly more −1 oocytes displaying a punctate morphology than does wild type. The punctate morphology maintained in *daf-2* oocytes suggests an alternative mechanism may be employed to combat age-related reproductive decline, distinct from *daf-2’s* maintenance of elongated mitochondria in aged somatic tissues.

### Fission and fusion are necessary for normal reproductive function with age

The punctate morphology of the mitochondria in *daf*-2 oocytes is distinct from their morphology in wild-type oocytes and somatic tissues, and also distinct from their morphology in *daf-2* somatic tissues, which suggests mitochondrial dynamics are differentially regulated in *daf-*2 reproduction. Therefore, we investigated the roles of regulators of mitochondrial dynamics in reproduction, DRP-1 (Dynamin-Related Protein 1) and FZO-1 (mitofusin 1/2). *C. elegans* mutants that lack fission (*drp-1(tm1108)*) or fusion (*fzo-1(tm1133)*) do not exhibit differences in lifespan (Supplementary Figures S2A and S2B; Weir et al., 2017; Yang et al., 2011), but the roles that fission and fusion play in reproductive span were still unknown. To test this, we measured their reproductive spans and assessed the oocyte quality of *drp-1* and *fzo-1* genetic null mutants. We confirmed that the mitochondria in *drp-1* mutant mature (−1) oocytes are elongated, while *fzo-1* mutant oocyte mitochondria are punctate (Figures 2A and 2B), as expected given their previously described functions.

**Figure 2.**
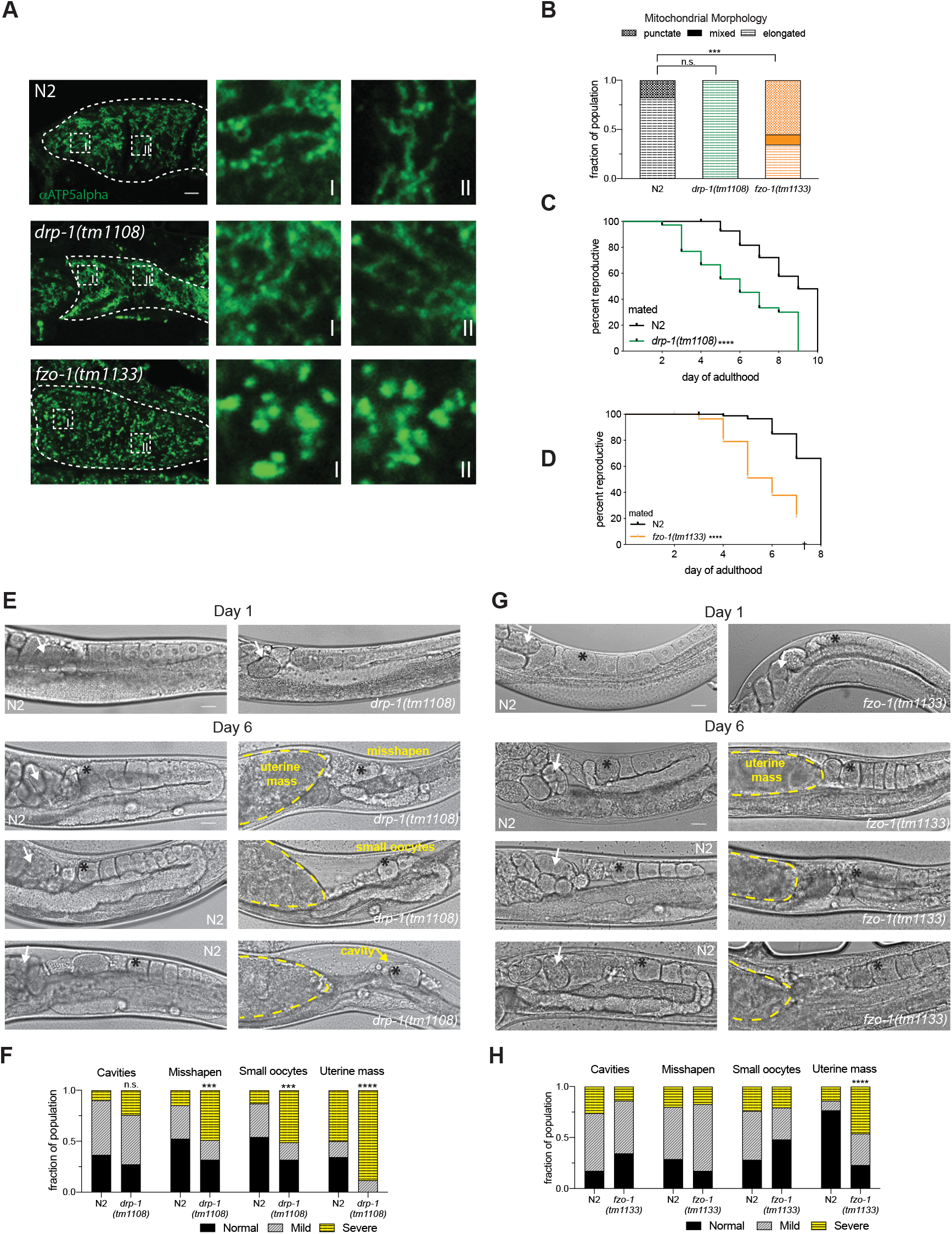
Fission and fusion are necessary for normal reproductive function with age. (A) Mitochondria morphology in mature oocytes of N2, *drp-1(tm1108)*, and *fzo-1(tm1133)*. Adult Day 3 germlines were dissected and stained with anti-ATP5α mitochondrial membrane marker. Representative images highlight the different morphologies in N2 (elongated), *drp-1(tm1108)* (elongated), and *fzo-1(tm1133)* (punctate). Scale bar 10um. White-dotted boxes are enlarged (right) of the morphology in the −1 oocyte (I) and the −2 oocyte (II). Images in A were processed using the same parameters across the entire image (i.e. adjusting for brightness and contrast). (B) Quantification results of mitochondrial morphology in the −1 oocyte in Day 3 adults of N2 (n=34), *drp-1(tm1108)* (n=29), and *fzo-1(tm1133)* (n=29). (C) *drp-1(tm1108)* has a reduced reproductive span in hermaphrodites (n=96-109) mated with males. (D) *fzo-1(tm1133)* has a reduced reproductive span in hermaphrodites (n=70-89) mated with males. (E) In aged adults (Day 6), *drp-1(tm1108)* have more misshapen, small oocytes and more uterine masses (signs of oocyte quality decline) compared to N2. Highlighted features include: embryos in the uterus (white arrows), the −1 oocyte (black asterisk), oocyte quality-decline phenotypes (yellow text and arrows), uterine mass (yellow dotted outline). (F) Quantification of oocyte quality decline phenotypes in N2(n=64) and *drp-1(tm1108)* (n=51). (G) In mated Day 6 adults, *fzo-1(tm1133)* has an increased number of hermaphrodites with a uterine mass instead of developing embryos. Highlighted features same as E. Scale bars 20um. (H) Quantification of oocyte quality decline phenotypes in N2(n=38) and *fzo-1(tm1133)* (n=34). *p ≤ 0.05, **p ≤ 0.01, ***p ≤ 0.001, ****p ≤ 0.0001.

Next, we asked what effect the disruption of mitochondrial dynamics has on reproductive span. To assess whether mitochondrial dynamics specifically play a role in oocyte quality determination of reproductive span, we performed a mated reproductive span assay with wild-type males to control for possible impairment of mutant sperm as well as depletion of self-sperm (Luo et al., 2010, 2009). We found that the mated reproductive spans of both *drp-1(tm1108)* (Figure 2C) and *fzo-1(tm1133)* (Figure 2D) are significantly shorter than wild type’s reproductive span, suggesting that defective oocytes are at least partly responsible for impaired reproduction of fission and fusion mutants, and that disruption in either fission or fusion is deleterious for maximal reproductive span.

To determine whether loss of fission or fusion affects oocyte quality, we imaged age-matched hermaphrodite germlines and assessed oocyte quality phenotypes. In Day 6 adults, a time point near the end of wild type’s reproductive span, there is a significantly higher incidence of misshapen and smaller oocytes in *drp-1(tm1108)* fission mutants compared to N2, and a large number of *drp-1(tm1108)* had a mass of oocytes or other cellular material in the uterus (termed “uterine mass”) rather than normally developing embryos (Figure 2E and 2F). Importantly, the oocyte quality phenotypes observed in aged *drp-1(tm1108)* are not present in young adults (Day 1, Figure 2F) suggesting this decline is dependent on age. Aged *fzo-1* fusion mutant oocytes display large uterine masses in place of developing embryos in the majority of the population (Figure 3G and 3H). Again, this uterine mass does not appear in young adults (Day 1, Figure 3H). These results suggest that the primary regulators of both fission and fusion are required for normal reproductive health through the maintenance of oocyte quality with age.

**Figure 3:**
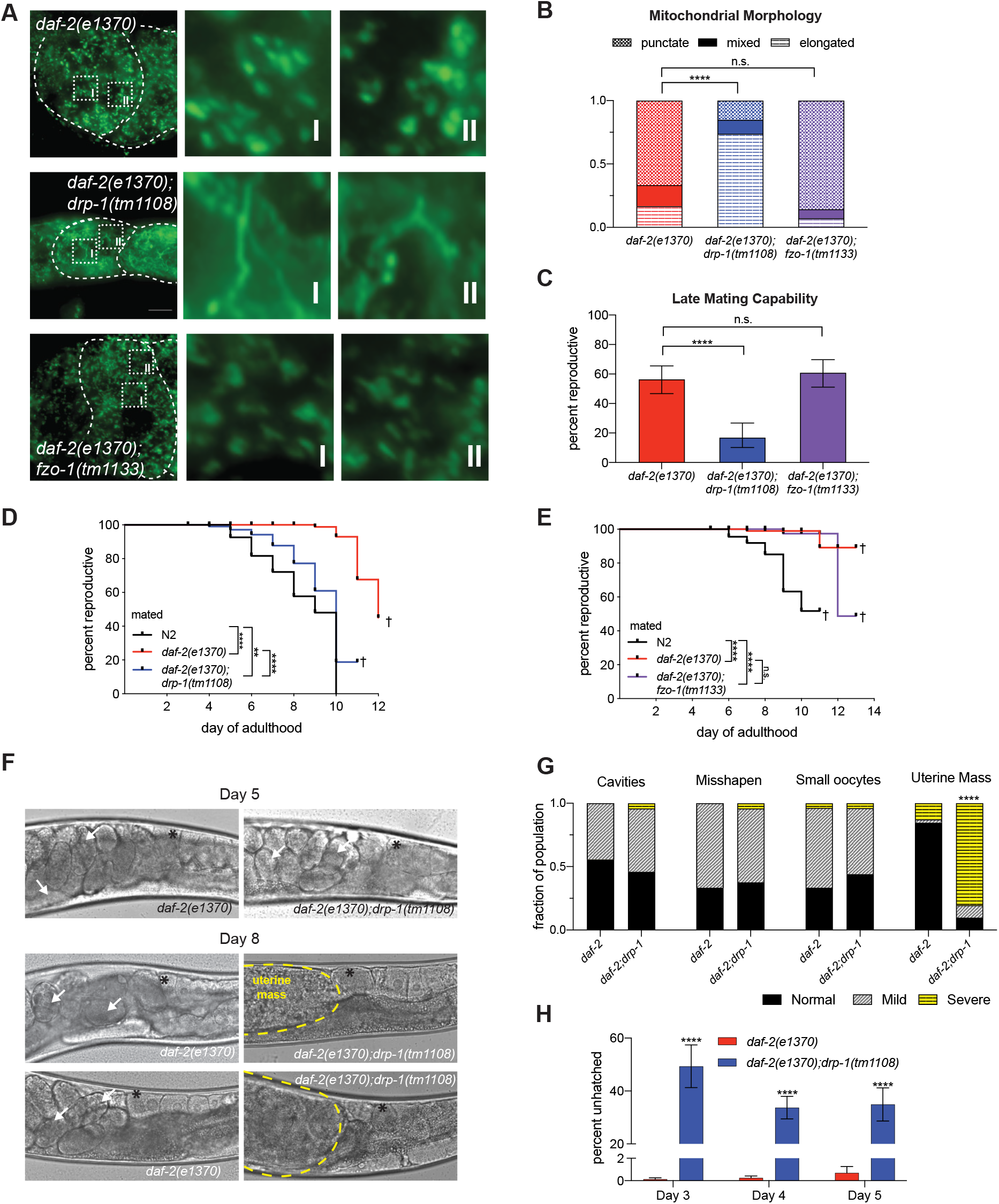
*daf-2* requires fission, not fusion, for extended reproductive function and oocyte quality maintenance with age. Representative images of Day 4 adult mitochondria morphology in the −1 oocyte using anti-ATP5α to mark mitochondria membranes. White-dotted boxes are enlarged (right) of the morphology in the −1 oocyte (I and II). Scale bar 10um. Images in A were processed using the same parameters across the entire image (i.e. adjusting for brightness and contrast). (B) Blind scoring results of mitochondria morphology of the −1 oocyte in three categories: punctate, mixed, or elongated. *daf-2(e1370);drp-1(tm1108)* (n=46) have majority elongated mitochondria, *daf-2(e1370);fzo-1(tm1133)* retains punctate morphology (n=56) similar to *daf-2(e1370)* (n=30). (C) *daf-2(e137)* (n=103) and *daf-2(e1370);fzo-1(tm1133)* (n=102) have high late-mating capability after aged Day 7 adults were mated with young *fog-2* males. *daf-2(e1370);drp-1(tm1108)* (n=79) are significantly less reproductive. (D) Fission knockout with *daf-2(e1370);drp-1(tm1108)* (n=96) has a reduced reproductive span compared to *daf-2(e1370)* (n=121) in mated hermaphrodites. (E) With fusion knockout, the reproductive span is still extended in *daf-2(e1370);fzo-1(tm1133)* (n=101) compared to *daf-2(e1370)* (n=102) of mated hermaphrodites. † indicates loss of hermaphrodites due to high rates of matricide. (F) Representative images of oocyte quality in aged adult (Day 8) germlines. *daf-2* has developing embryos in the uterus (white arrows) while *daf-2;drp-1* has uterine masses (yellow dashed line). There are still developing embryos in *daf-2;drp-1* in younger adults (upper panel). Black asterisks mark the −1 oocyte. Scale bar is 20um. (G) Quantification of oocyte quality phenotypes from *daf-2(e1370)* (n=41) and *daf-2(e1370);drp-1(tm1108)* (n=39). (H) *daf-2(e1370);drp-1(tm1108)* has a high fraction of unhatched embryos compared to *daf-2(e1370)*. Every 24 hours plates were counted for hatched progeny, unhatched embryos, and unfertilized oocytes. Representative data from days 3, 4, and 5 of adulthood are shown due to high rates of matricide after day 5 (n=11-13). **p ≤ 0.01, ****p ≤ 0.0001.

### *daf-2* requires fission, not fusion, for extended reproductive function and oocyte quality maintenance with age

*daf-2* oocyte mitochondrial morphology is punctate rather than elongated, and this morphology is retained throughout reproduction. We next asked whether this punctate state is necessary for *daf-2*‘s extended reproductive span. If *daf-2* specifically requires the punctate morphology in oocytes, then eliminating fission should prevent *daf-2’s* maintenance of reproduction. To test whether fission is required for *daf-2’s* extended reproduction, we crossed *daf-2(e1370)* with *drp-1(tm1108),* and first confirmed that the mitochondria in the −1 oocyte are elongated, not punctate (Figure 3A and 3B). We then tested their reproduction and found that *daf-2(e1370);drp-1(tm1108)* mutants have a significantly reduced late-mating capability (Figure 3C), and the reproductive span extension of *daf-*2 is greatly reduced (Figure 3D). By contrast, promoting fission by knocking out fusion protein FZO-1 does not affect reproduction: like *daf-2*, the −1 oocytes of *daf-2(e1370);fzo-1(tm1133)* mutants exhibit primarily punctate mitochondria (Figure 3A and 3B). Similarly, loss of FZO-1 does not affect *daf-2’s* reproductive capability in aged hermaphrodites (Figure 3C) or extended reproductive span (Figure 3E). Therefore, *daf-2* requires fission, not fusion, to preserve reproductive function with age.

We next asked whether the requirement of DRP-1 for reproductive longevity in *daf*-2 is through the maintenance of oocyte quality. We found the majority of mated Day 8 *daf-2(e1370);drp-1(tm1108)* mutants have large masses, instead of developing embryos, in their uteri (Figures 3F and 3G). The “uterine mass” phenotype is a sign of oocyte quality decline and is not present in younger adults (Day 5, Figure 3G). Consistent with a loss of oocyte quality, *daf-2(e1370);drp-1(tm1108)* mutants lay a high percentage of unhatched embryos on days 3, 4, and 5 of adulthood (Figure 3H). Together, our data suggest that retaining the punctate state of mitochondria through fission in oocytes is required for *daf-2’s* maintenance of high-quality oocytes with age, which is critical for its extended reproductive span.

### Mitophagy is required for *daf-2*’s reproductive longevity

One possible benefit to *daf-*2 maintaining punctate mitochondria in oocytes is they may be primed for mitophagy, which provides quality control for damaged mitochondria. Mitochondria that have accumulated damage divide asymmetrically through fission (Twig et al., 2008) and activate mitophagy to metabolize mitochondria that are beyond repair (Youle and van der Bliek, 2012). The main mitophagy pathway regulator in *C. elegans* is PINK-1 (PTEN induced kinase 1) (Pickrell and Youle, 2015); to test its role, we examined the phenotypes of *daf-2(e1370);pink-1(tm1133)* double mutants. Aged (Day 7) *daf-2(e1370);pink-1(tm1133)* double mutants have reduced reproductive capability after late mating (Figure 4A). There is no difference between the reproductive spans of N2 and *pink-1(tm1133)* (Figure 4B), suggesting that PINK-1 may be specifically required for reproductive longevity mechanisms that favor the elimination of damaged, fissioned mitochondria.

**Figure 4:**
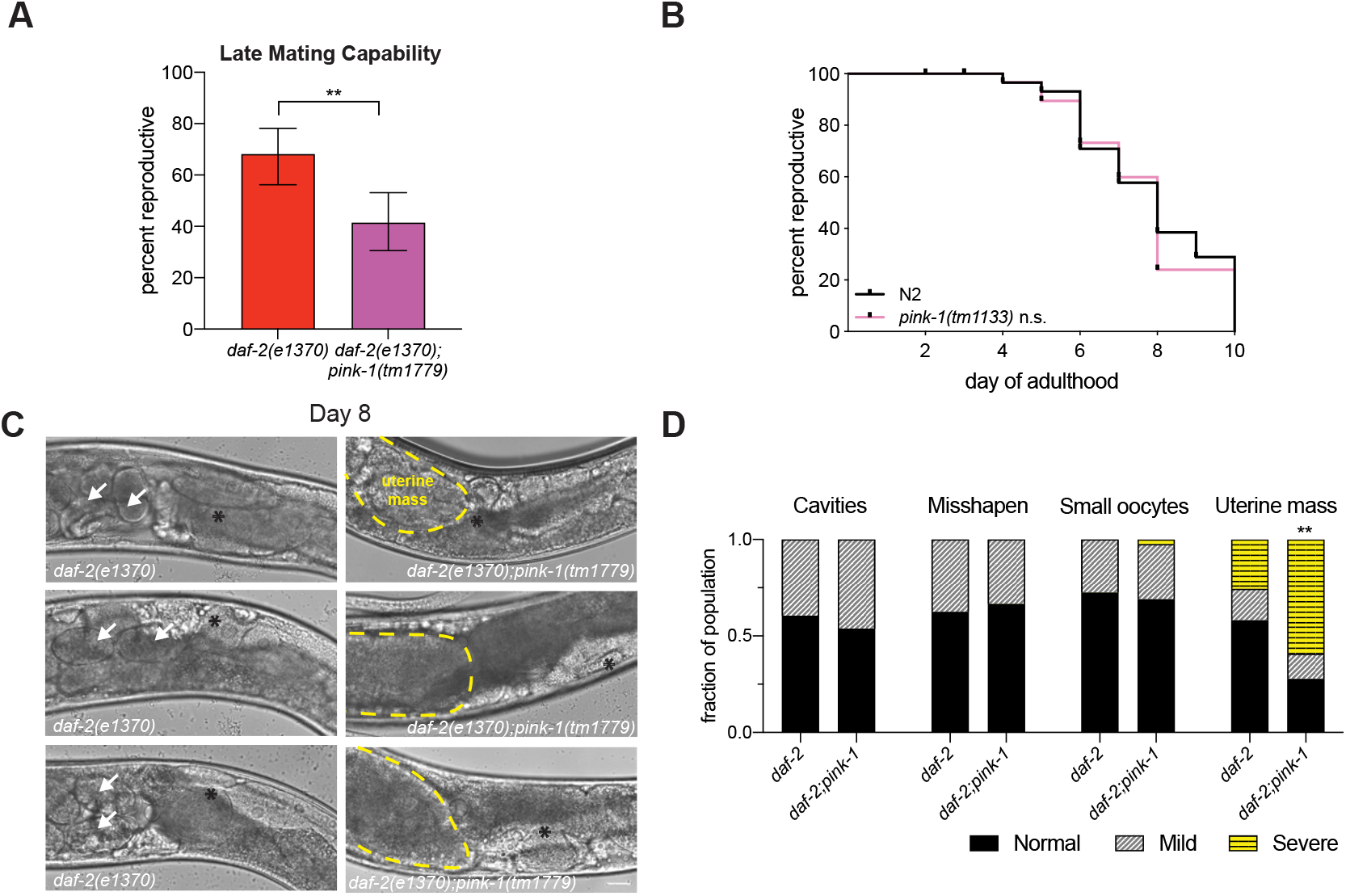
Mitophagy is required for *daf-2*’s extended reproductive capability due to loss of oocyte quality with age. Knocking down mitophagy in *daf-2(e1370);pink-1(tm1779)* (n=66) reduces late-mating capability compared to *daf-2(e1370)* (n=70). Late-mating performed on age matched Day 7 hermaphrodites provided with young males. Error bars represent 95% confidence interval. (B) PINK-1 mediated mitophagy is not required for normal reproductive span. There is no significant difference between the mated reproductive spans of N2 (n=59) and *pink-1(tm1133)* (n=63). (C) Loss of oocyte quality in *daf-2(e1370);pink-1(tm1779)* in aged adult germlines (Day 8) is due to a “uterine mass” phenotype in place of developing embryos. Scale bar is 20um. (D) Quantification of oocyte quality decline phenotypes in *daf-2(e1370)* (n=45) and *daf-2(e1370);drp-1(tm1108)* (n=51). **p ≤ 0.01, ****p ≤ 0.0001.

Similar to *daf-2(e1370);drp-1(tm1108)* mutants, mated Day 8 *daf-2;pink-1* hermaphrodites exhibited a significant number of uterine masses rather than developing embryos (Figure 4C-D; compare to 3F-G), suggesting that the uterine mass phenotype is a result of mitochondrial dysfunction in aging oocytes brought on by the loss of mitophagy. Taken together, our results support a model in which punctate mitochondria are primed for mitophagy in *daf-2* oocytes to promote reproductive longevity.

### Fission and mitophagy are required for *daf-*2’s reproductive span, not somatic lifespan

Next we asked whether mitochondrial morphology and mitophagy regulators also play a role in *daf-2’s* somatic longevity. However, we found that loss of *drp-1*, *fzo-1*, and *pink-1* each had no effect on the extended lifespan of *daf-2* mutants (Figure 5A-C), suggesting that the mechanism we have identified involving mitochondrial dynamics and mitophagy is specific to reproductive longevity through oocyte quality maintenance with age.

**Figure 5:**
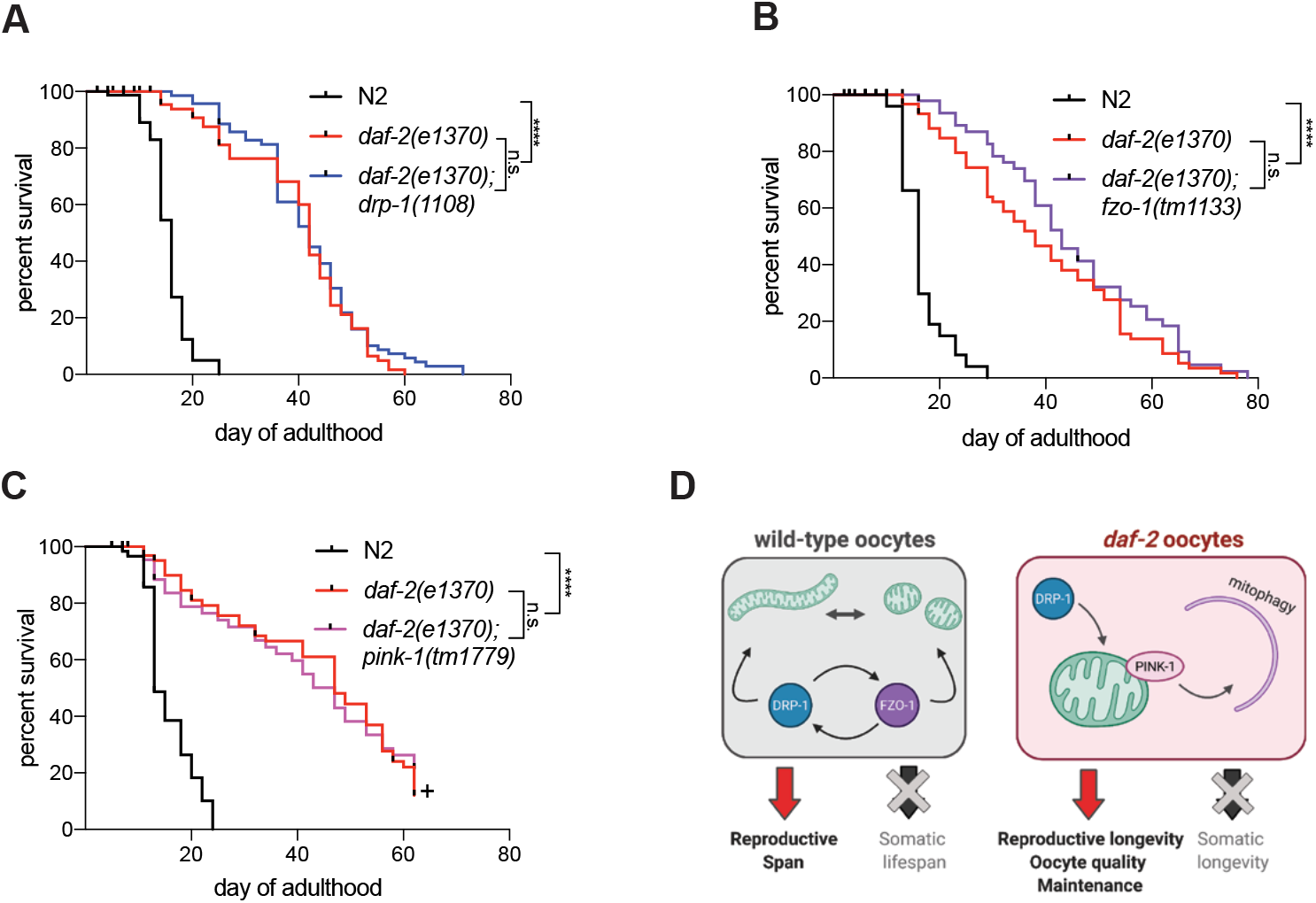
Fission and mitophagy are required for *daf-2*’s reproductive span, not somatic lifespan. (A) *daf-2(e1370);drp-1(tm1108)* (n=81) has an extended lifespan like *daf-2(e1370)* alone (n=80). (B) *daf-2(e1370);fzo-1(tm1133)* (n=80) has an extended lifespan comparable to *daf-2(e1370)* (n=81). (C) There is no change between the lifespans of *daf-2(e1370);pink-1(tm1779)* compared to *daf-2(e1370)* alone (n=66-79). + indicates censorship due to covid-19 shutdown. (D) Wild type requires both fission and fusion, but not mitophagy, to maintain a normal reproductive span. Reproductive longevity in *daf-2* relies on fissioned mitochondria, which are then primed for and utilize mitophagy to maintain reproductive longevity through oocyte quality maintenance. ****p ≤ 0.0001.

## Discussion

Here we sought to determine whether mitochondrial morphology influences oocyte quality and reproductive span, and whether there is an ideal morphology that promotes reproductive health. There appear to be two distinct mechanisms involving mitochondrial dynamics during reproductive aging. Wild-type oocytes require both fission and fusion to maintain normal reproductive function with age, suggesting that this mitochondrial dynamics are required to support normal reproductive function in wild-type animals (Figure 5D).

*daf-2* appears to use a different strategy to promote reproductive longevity, requiring DRP-1 to tip the balance of mitochondria to the fissioned, punctate state. We found no evidence that *daf-2* oocyte mitochondria transition into a tubular, elongated state. This result was surprising, as punctate mitochondria are usually thought to be dysfunctional, and are often referred to as “fragmented.” However, in the case of *daf-2’s* reproduction, punctate mitochondria are beneficial. This benefit may be due, in part, to the fact that fissioned mitochondria activate the mitophagy pathway (Youle and van der Bliek, 2012); indeed, we find that mitophagy is required for *daf-*2’s extended reproductive span. In this model, dysfunctional mitochondria are removed from the mitochondria pool through PINK-1-mediated mitophagy, while healthy mitochondria are retained in *daf-2* oocytes. The healthy population of mitochondria in oocytes is maintained with age and assists in slowing oocyte quality decline, thereby extending reproduction (Figure 5D).

Insulin/IGF-1 longevity mutants retain elongated mitochondria in muscle and neurons later in life (Jiang et al., 2015; Lamarche et al., 2018; Morsci et al., 2016; Regmi et al., 2014), which supports the hypothesis that fusion promotes longevity. However, promoting fusion, by knocking out the fission protein DRP-1, or by overexpressing the fusion protein FZO-1, has no effect on lifespan (Weir et al., 2017); Supplemental Figure 2). This data suggest that neither fusion nor fission is the main driver of somatic lifespan. However, we have shown here that mitochondrial morphology and dynamics specifically regulate *C. elegans* reproductive span. The requirement of mitochondrial dynamics specifically for reproduction may be due, in part, to the fact that the germline contains the vast majority (approximately 85%) of mitochondria in the *C. elegans* adult, and up to a 6-fold increase in adult mtDNA copy number may be attributed to oocyte production (Tsang and Lemire, 2002). The sheer abundance of mitochondria in the germline, coupled to the importance of healthy progeny, amplifies the need for high-quality mitochondria during reproduction.

The tissue-specific effect that mitochondrial dynamics has on reproduction is also mirrored in *daf-2*. The loss of *drp-1* and *fzo-1* from the *daf-2(e1370)* background has no effect on lifespan (Figure 5A-B). PINK-1-mediated mitophagy also has no role in *daf-2* somatic longevity (Figure 5C), although autophagic activity (Meléndez et al., 2003; Palikaras et al., 2015), mitophagy, and mitochondrial biogenesis mechanisms are upregulated in *daf-2* (Palikaras et al., 2015). These results suggest that somatic lifespan and reproductive span have distinct requirements for mitochondrial morphology and employ different mechanisms to combat age-related mitochondrial dysfunction. A PINK-1-mediated mitophagy mechanism found in *daf-2* is an alternative strategy to promote reproductive longevity, distinct from the assumption that tubular, elongated mitochondria are healthier. Drosophila uses a similar mechanism in early oogenesis, in which mitochondria undergo fragmentation to selectively remove mitochondria with mutated mtDNA through an Atg8 and BNIP3 mitophagy/autophagy pathway (Lieber et al., 2019).

Mechanisms to slow reproductive decline have become increasingly important as the average maternal age increases. Our results demonstrate that studying longevity mutants, like the IIS longevity mutant *daf-2*, is a viable approach to uncover unique mechanisms to combat age-related reproductive decline, and may provide insight into new strategies to promote reproductive longevity. The requirement for mitophagy in *daf-2* to promote reproductive span extension provides an alternative strategy to combat age-related reproductive decline and supports investigating mitophagy as a means to promote reproductive health with age.

## Conflict of Interest

The authors declare no competing interests.

## Acknowledgements

We thank the *Caenorhabditis* Genetics Center (CGC) and the National BioResource Project (NBRP) for strains, Jasmin Ashraf, Will Keyes, and Rebecca S. Moore for help with experiments, members of the Murphy Lab for discussion and feedback on the manuscript, Biorender.com for graphical abstract and model figure design software. Imaging was performed with support from the Confocal Imaging Facility, a Nikon Center of Excellence, in the Department of Molecular Biology at Princeton University. C.T.M. is the Director of the Glenn Center for Aging Research at Princeton and an HHMI-Simons Faculty Scholar. V.C. was supported by T32GM007388 (NGMIS). This project was funded in part by grant number GCRLE-0220 from the *Global Consortium for Reproductive Longevity and Equality* through the Buck Institute, made possible by the Bia-Echo Foundation.

---

**Figure S1:**
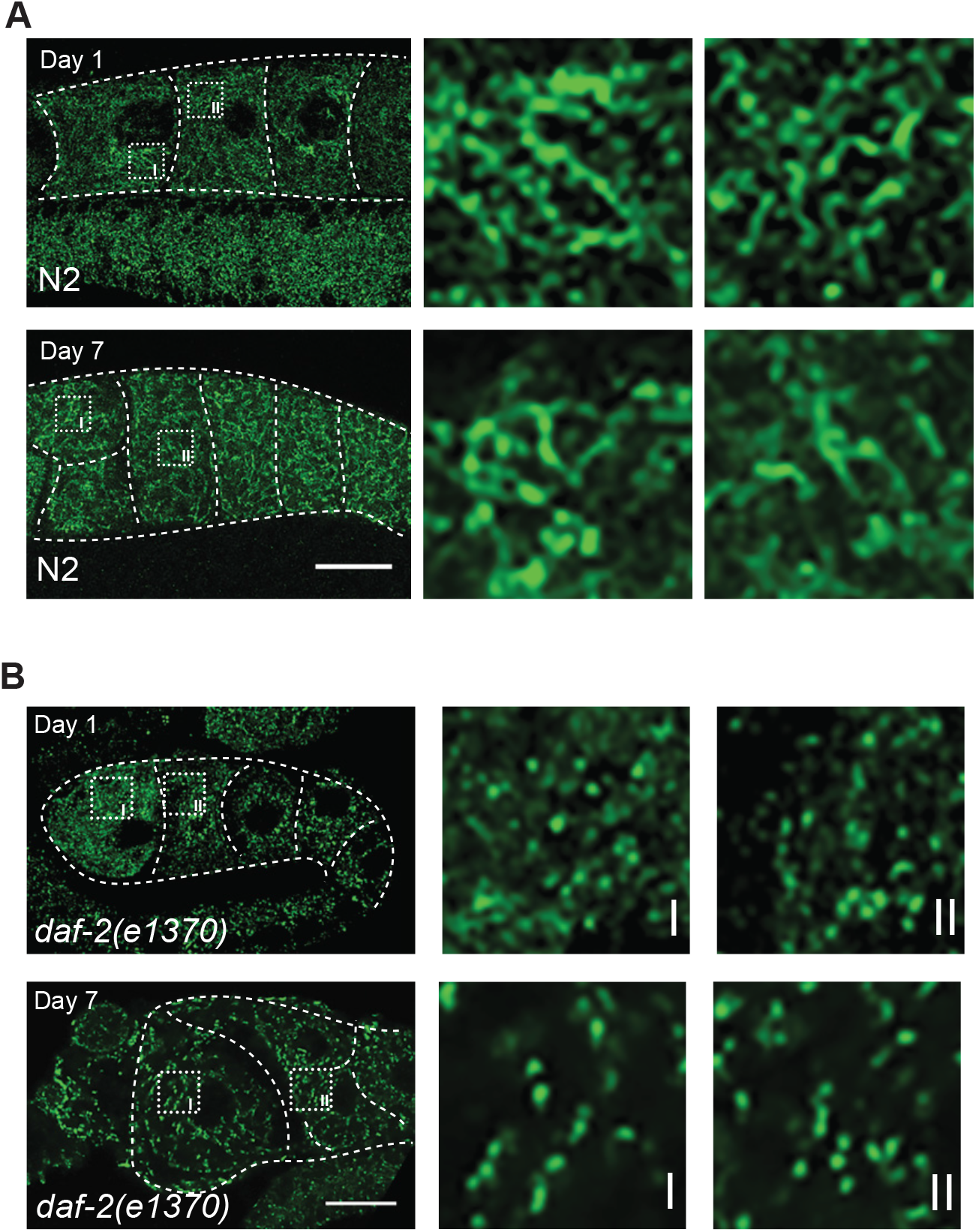
Mitochondria morphology does not change with age in N2 or *daf-2(e1370)* Morphology images of N2 Day 1 and Day 7 (A) and *daf-2(e1370)* (B). Mitochondria are stained with ATP5α to mark membranes.

**Figure S2:**
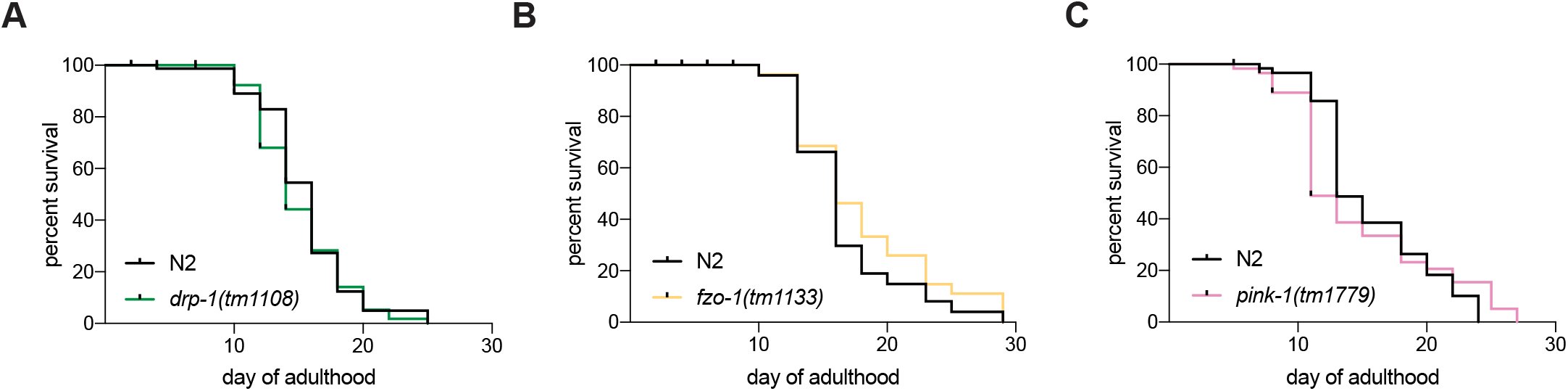
Mitochondrial dynamics are specifically required for reproduction. (A) Lifespans are not affected by loss of *drp-1(tm1108)* (n=80-81), (B) *fzo-1(tm1133)* (n=80-83) or *pink-1(tm1779)* (n=62-69).

## METHODS

### General worm maintenance

The following strains were used in this study: N2 Bristol strain as wild type worms, *fog-2(q71) V* males for mating experiments, *daf-2(e1370)*, *drp-1(tm1108)*, *fzo-1(tm1133)*, *pink-1(tm1179)*. The following crosses were performed: *daf-2(e1370);drp-1(tm1108)*, *daf-2(e1370);fzo-1(tm1133)*, and *daf-2(e1370);pink-1(tm1170)*.

All strains were cultured using standard methods (Brenner, 1974). For all experiments worms were maintained at 20C. Nematode growth medium (NGM: 3 g/L NaCl, 2.5 g/L Bacto-peptone, 17 g/L Bacto-agar in distilled water, with 1mL/L cholesterol (5 mg/mL in ethanol), 1 mL/L 1M CaCl_2_, 1 mL/L 1M MgSO_4_, and 25 mL/L 1M potassium phosphate buffer (pH 6.0) added to molten agar after autoclaving. Plates were seeded with OP50 *E. coli* for *ad libitum* feeding. To synchronize experimental groups, gravid hermaphrodites were used to collect eggs by exposing them to a 15% hypochlorite solution (e.g., 8.0 mL water, 0.5 mL 5N KOH, 1.5 mL sodium hypochlorite), followed by repeated washing of collected eggs in M9 buffer (Brenner, 1974).

### Reproductive span and Lifespan Assays

Mated reproductive spans were performed as previously described (Luo et al., 2009). Worms were synchronized with a hypochlorite solution and selected for experiments at the L4 stage. Hermaphrodites were mated with *fog-2(q31)* males at a 2:1 ratio for 24 hrs from L4 to Day 1 of adulthood. Each hermaphrodite was then singled onto 35mm NGM plates and moved to new plates every 24 hours until the end of the reproductive span. 48 hours after removal plates were screened for progeny to confirm reproductive span, including male progeny to confirm a successful mating. Reproductive cessation was defined as the last day of progeny production preceding two full days without progeny. Hermaphrodites were censored on the day of matricide (defined as progeny hatched within mother). All reproductive spans were performed at 20C.

Lifespan assays were performed as previously described (Kenyon et al., 1993). In brief, groups of hypochlorite-synchronized hermaphrodites (<15 per plate) were placed on plates at the L4 stage. The hermaphrodites were transferred to freshly seeded plates every two days when producing progeny, and 2-3 days thereafter. Worms were censored on the day of matricide (defined as progeny hatching within mother), abnormal vulva structures, or loss. Worms were defined as dead when they no longer responded to touch. All lifespans were performed at 20°C.

### Oocyte quality assays

Assays were performed as previously described (Luo et al., 2010; Templeman et al., 2018). Hypochlorite-synchronized hermaphrodites were mated on either L4 or Day 1 of adulthood. Mating was confirmed as in the reproductive span by confirming the presence of male progeny. Groups of hermaphrodites were maintained until day of experiment (Day 6-Day 8 of adulthood). On day of experiment, hermaphrodites were added to slides with 3-4% agarose pads in M9 with levamisole. DIC images were captured on a Nikon eclipse Ti at 60x magnification. To perform scoring, scorer was blinded to genotype and each image was given a score (normal, mild, severe) in each oocyte quality category. Note that in experiments using *daf-2(e1370)*, worms were added to plates with serotonin for 1 hour on Day 6 and Day 8 to purge embryos and prevent matricide due to bagging.

### Embryonic Lethality Assay

Assay was performed as described (Luo et al., 2010). L4 hermaphrodites were singled onto 35mm NGM plates and males were added at a 3:1 ratio for mating. Mating was confirmed as in the reproductive span. Every 24 hours hermaphrodites were moved to new plates. Approximately 24 hours after removal, plates were scored for numbers of progeny, unhatched embryos, and unfertilized oocytes. Counting continued throughout reproductive span until the hermaphrodites’ numbers fell below 10. Percent unhatched is the proportion of embryos that failed to hatch after 24 hours.

### Late-mating assay

Late-mating assays were performed as previously described (Templeman et al., 2020). Hypochlorite-synchronized worms were selected at the L4 stage. Hermaphrodites were aged to Day 7, moving every two days to separate from progeny. On day of late-mating, hermaphrodites were individually placed onto 35mm plates seeded with 25uL spots of OP50. *fog-2(q71)* males were added at a 3:1 ratio and presence of progeny was scored 4-5 days after males were added to the plates.

### Mitochondria morphology assay

Mitochondria morphology was assessed using immunohistochemistry staining with the mitochondria membrane marker ATP5α. Germline immunostaining was performed using the protocol as in (Shaham, 2006). In brief, hypochlorite-synchronized hermaphrodites were maintained until the day of interest. Hermaphrodite germlines were dissected and fixed using a methanol/acetone fixation. Primary antibody from AbCam Mouse Anti-ATP5α antibody [15H4C4]. Invitrogen Goat Polyclonal Secondary Antibody Alexa Fluor™ Plus 555. Imaging was performed on point scanning confocal Nikon A1 at either 100x or 60x or Nikon eclipse Ti. Images were taken on a Z-stack (0.7-1.0 um steps) focusing on the −1 oocyte. For mitochondria morphology quantification, images from 2-4 genotypes were collected and file names were blinded. Each Z-stack was then assessed for a morphology qualification (punctate, mixed, elongated). Images were processed in FIJI and Adobe Photoshop. Each figure of images shown were from the same experiment, imaged on the same microscope at the same magnification, and with the same exposure. All images in each figure were processed in the same way across the entire image (e.g. adjusting for brightness and contrast).

## QUANTIFICATION AND STATISTICAL ANALYSIS

Lifespans and reproductive span assays were assessed using the standard Kaplan-Meier log rank survival tests. The first day of adulthood was defined as t=0. Oocyte quality assays used a chi square test to determine whether there was any significant difference between populations for each oocyte quality category. Embryonic lethality assays used a two-way Anova to compare across experimental days and genotypes. In late-mating assays chi-square tests were used to compare percentages of reproductive and non-reproductive populations across genotypes. For morphology analysis, chi-square tests were used to compare the percentages of the population within each morphology category.

All experiments were repeated on separate days with separate, independent populations. Details of each represented experiment, including sample sizes, can be found in the figure legend. All figures in the Article and Supplement shown are of 1/3 biological replicates with exception of the mitochondrial morphology scoring, which are pooled data from independent experiments. Prism 9 software was used for statistical analysis.

## Notes

### Competing Interest Statement

The authors have declared no competing interest.

